# Attractor-like spatial representations in the Subicular Complex network

**DOI:** 10.1101/2020.02.05.935379

**Authors:** Apoorv Sharma, Indrajith R. Nair, Yoganarasimha Doreswamy

## Abstract

Distinct computations are performed at multiple brain regions during encoding of the spatial environments. Neural representations in the hippocampal, entorhinal and head direction (HD) networks during spatial navigation have been clearly documented, while the representational properties of the Subicular Complex (SC) network is rather unexplored, even though it has extensive anatomical connections with various brain regions involved in spatial information processing. Here, we report a global cue controlled highly coherent representation of the cue-conflict environment in the SC network, along with strong coupling between HD cells and Spatial cells. We propose that the attractor dynamics in the SC network might play a critical role in orientation of the spatial representations, thus providing a “reference map” of the environment for further processing at other networks.

## Introduction

The Subicular Complex (SC), consisting of the Subiculum (SUB), Presubiculum (PrS) and Parasubiculum (PaS) regions, is strategically located between the hippocampus and the entorhinal cortex (EC), and has extensive anatomical connections with various cortical and sub-cortical areas of the brain (*1*). Previous studies have reported the spatial and directional correlates of neuronal firing in the SC, such as the place cells (*2-5*), head direction (HD) cells (*6, 7*), border cells (*8*) and grid cells (*7*), and their response to environmental manipulations (*9-12*).

The hippocampal and parahippocampal networks play a critical role during spatial navigation by encoding the salient features of the environment. In dynamically changing environments, distinct neural representations have been reported in specific regions of this network because of their intrinsic connectivity and anatomical inputs to them. The hippocampal place cell representations are governed by both the global and local cues of the environment (*13-15*), while a clear dissociation of global vs. local cue control of neural representations has been reported in the EC (*16*). In contrast, the anterior thalamus (ADN) HD network display global cue controlled highly coherent representations (*10, 15*). However, the nature of representation of space in the subicular complex (SC) network under these conditions still remain unclear.

## Results

To understand the representational properties of the SC in dynamically changing environments we conducted *in vivo* neurophysiological study by simultaneously recording the neural activity from all regions of the SC. Figure 1 A to C shows representative examples of tetrode localizations in SUB, PrS and PaS, respectively. In most of the experimental days, the cells were simultaneously recorded from a minimum of two regions of the SC (Fig. 1 D). Each day after the last recording session on track, a circular platform was placed on top of the track in the same room and a single recording session of 10 min duration was carried out to identify the cell type. Mean vector length, grid score and border score was calculated for each cell (Fig. 1 E) and categorised as either a border cell, grid cell or a head direction (HD) cell based on the threshold value obtained through data shuffling (Fig. 1 F). The remaining cells were classified as place cells provided their spatial information score was found to be significant (*p* < 0.05) in any of the track sessions recorded. Representative examples of a place, border, grid cell (Spatial cells) and HD cell based on this categorisation is shown in Fig.1 G to J.

**Figure 1.**
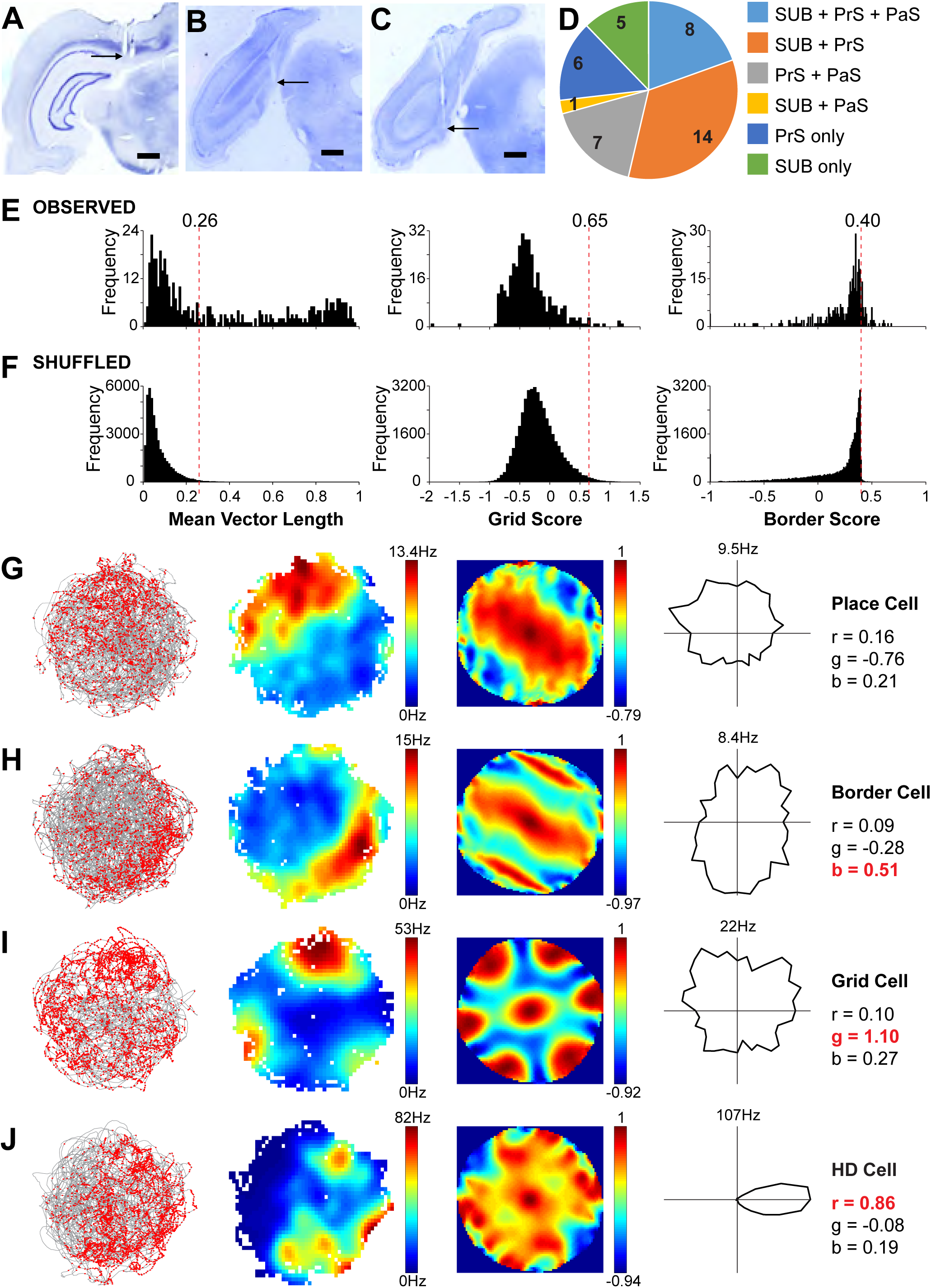
Neurophysiological recordings from the SC. **(A-C)** Representative examples of nissl-stained coronal sections of the rat brain showing tetrode tracks and recording sites (indicated by arrows) in SUB **(A)**, PrS **(B)** and PaS **(C)** regions of the SC. **D.** Pie chart showing the number of days cells were recorded from different regions of the SC simultaneously. **(E-F)** Distribution of mean vector length, grid score and border score values of all the cells recorded (**observed**) from different regions of the SC and distribution of these values after random shuffling of spike sequence (**shuffled**). Threshold values (99^th^ percentile of the shuffled data) are indicated by a red line. **(G-J)** Representative example each for different types of cells recorded from the SC region. Each row shows the trajectory of the rat (grey lines) superimposed with the spikes (red dots), firing rate map, spatial autocorrelogram, firing rate as a function of head direction, followed by their mean vector length (r), grid score (g) and border score (b) values. The scores that were above the threshold value are indicated in red.

We have recorded a total of 338 cells in different regions of the SC from 5 rats while they ran clock-wise (CW) on a textured circular track (the local cues) in presence of the global cues following a well-established experimental paradigm (*13-17*) (Fig. 2 A). Representative examples of HD cell tuning curves, firing patterns of spatial cells recorded from different rats across standard (STD) and mismatch (MIS) sessions within a day are shown in Fig. 2 B. All the cells maintained their preferred firing direction/location across different STD sessions, indicating the specificity and stability of the SC representation in familiar environments. In cue conflict conditions (MIS 180° and MIS 90°), the preferred firing direction of HD cells and the firing locations of spatial cells coherently rotated CW to an angle equal to the amount of rotation of the global cues.

**Figure 2.**
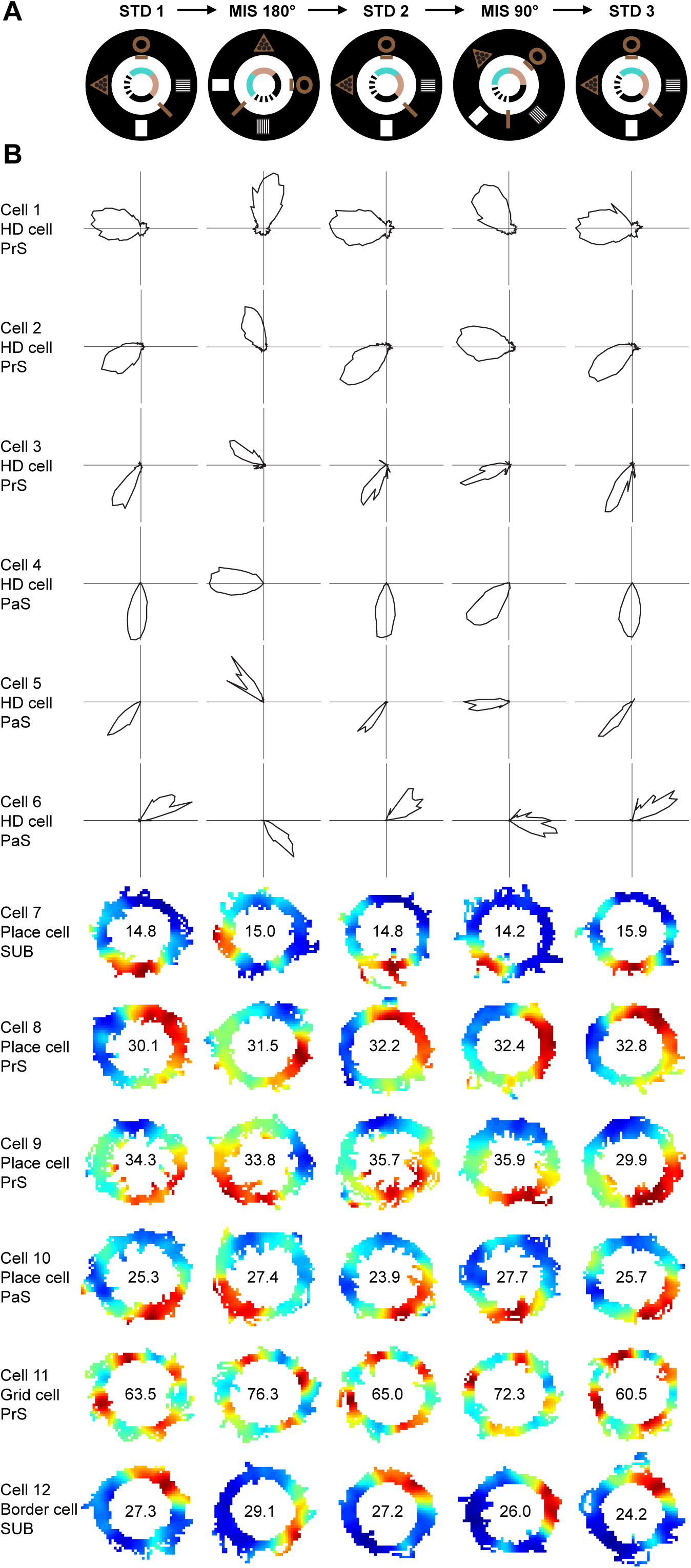
Experimental design and SC cell response to cue manipulations. **(A)** Schematic representation of the Local-global cue conflict experimental paradigm. **(B)** Representative examples of HD cells and Spatial cells recorded from different rats across STD and MIS sessions within a day. The axes for HD cells (1-6) are scaled for their maximum firing rates (112.6, 85.5, 17.0, 164.8, 22.4 and 17.2 Hz respectively). The numbers inside the firing rate maps indicate peak firing rate in Hz. The rate maps are color coded, with red color indicating >90% of the peak firing rate, blue no firing, and the successive decrements in peak firing rates are shown with intervening colors of the spectrum.

We quantified the amount of rotation of the cells in STD vs. STD and STD vs. MIS sessions by measuring the Pearson’s product-moment correlation of firing rate arrays. The angle at which highest correlation was observed was taken as the amount of rotation of preferred firing direction of a HD cell or firing field of spatial cells (*16*) and circular statistical analysis was conducted to calculate the mean angle of rotation of the population (angle of the mean vector) and variance of the distribution around the mean (length of the mean vector). Figure 3 shows the amount of rotation of the HD cells and the spatial cells recorded from different regions of the SC from all the rats (Fig. 3 A) and with all the regions combined (Fig. 3 B). The angle of the mean vector did not deviate significantly from zero between STD sessions suggesting no change in the preferred firing direction or firing fields, while deviated significantly from zero to an angle equal to the amount of rotation of the global cues in MIS sessions. Further, the length of the mean vector in all comparisons suggested that the angles were significantly clustered around the mean angle, in both the HD cells and the spatial cells. We observed no inter-regional differences in the pattern of representation when the responses from SUB, PrS and PaS regions were analysed separately (Fig. 3 A) and the representations in both the HD cells and the spatial cells of the SC were similar and highly coherent (Fig. 3 B). We further supplemented the above described categorical analysis by measuring the population responses to conflicting cue conditions. As there was no difference between HD cells and spatial cells representation, all the cells were grouped together and population responses were measured by creating 2D spatial correlation matrices from population firing rate vectors at each location on the track and then reduced to 1D polar plots for quantification (Fig. 4). In STD vs STD correlation matrices, the band of high correlation was observed on the central diagonal indicating no shift in firing direction/fields, while in STD vs MIS correlation matrices, the band of high correlation shifted CW corresponding to the amount of rotation of the global cues similarly in SUB, PrS and PaS (Fig. 4 A-C). The population response was stable across different STD sessions and rotated as an ensemble in global-local cue conflict conditions, suggesting a highly coherent network activity in the SC (Fig. 4 D).

**Figure 3.**
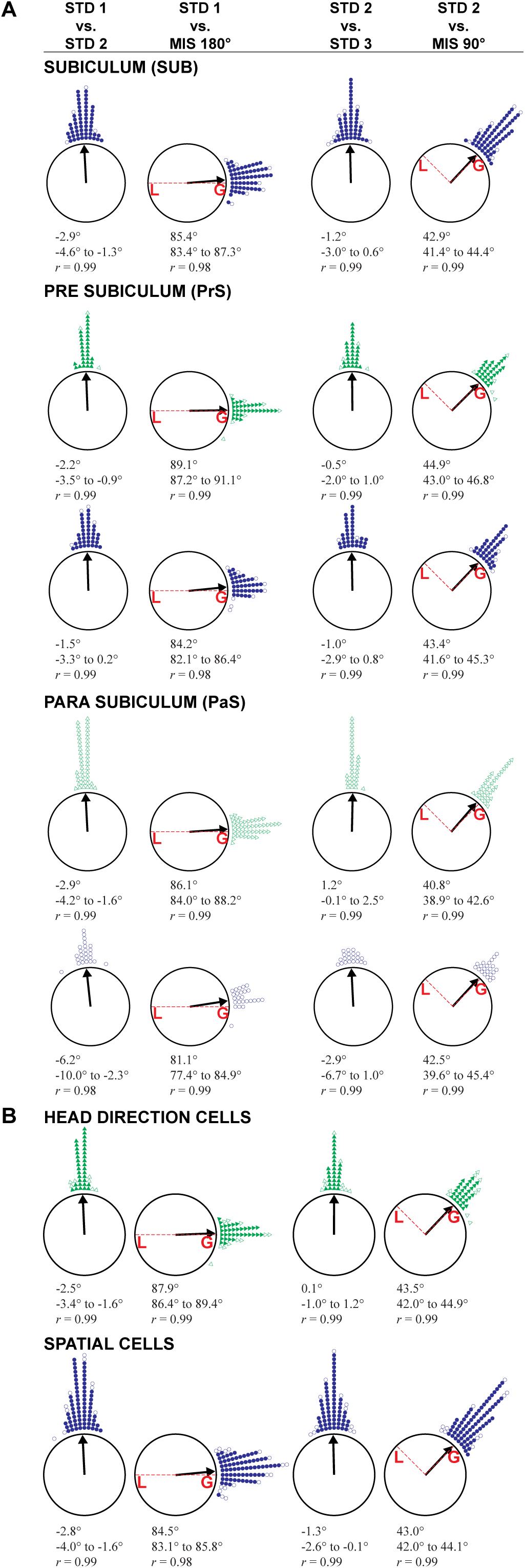
Global cue controlled highly coherent representation in the SC. **(A)** The amount of rotation of cells recorded from different regions of the SC between STDs and STD vs. MIS sessions is represented around the circle. (Green triangle = HD cells; Blue circle = Spatial cells; triangle/circle, open = 1 cell, filled = 2 cells). **(B)** The amount of rotation of HD cells and Spatial cells from all regions of the SC. (triangle/circle, open = 1 cell, filled = 3 cells). Values below each plot indicates the angle of mean vector, 95% confidence interval and length of the mean vector (*r*). All the values are significant at *p* < 0.0001. Red lines indicate the rotation angle of the local (L) and global (G) cues in MIS sessions.

**Figure 4.**
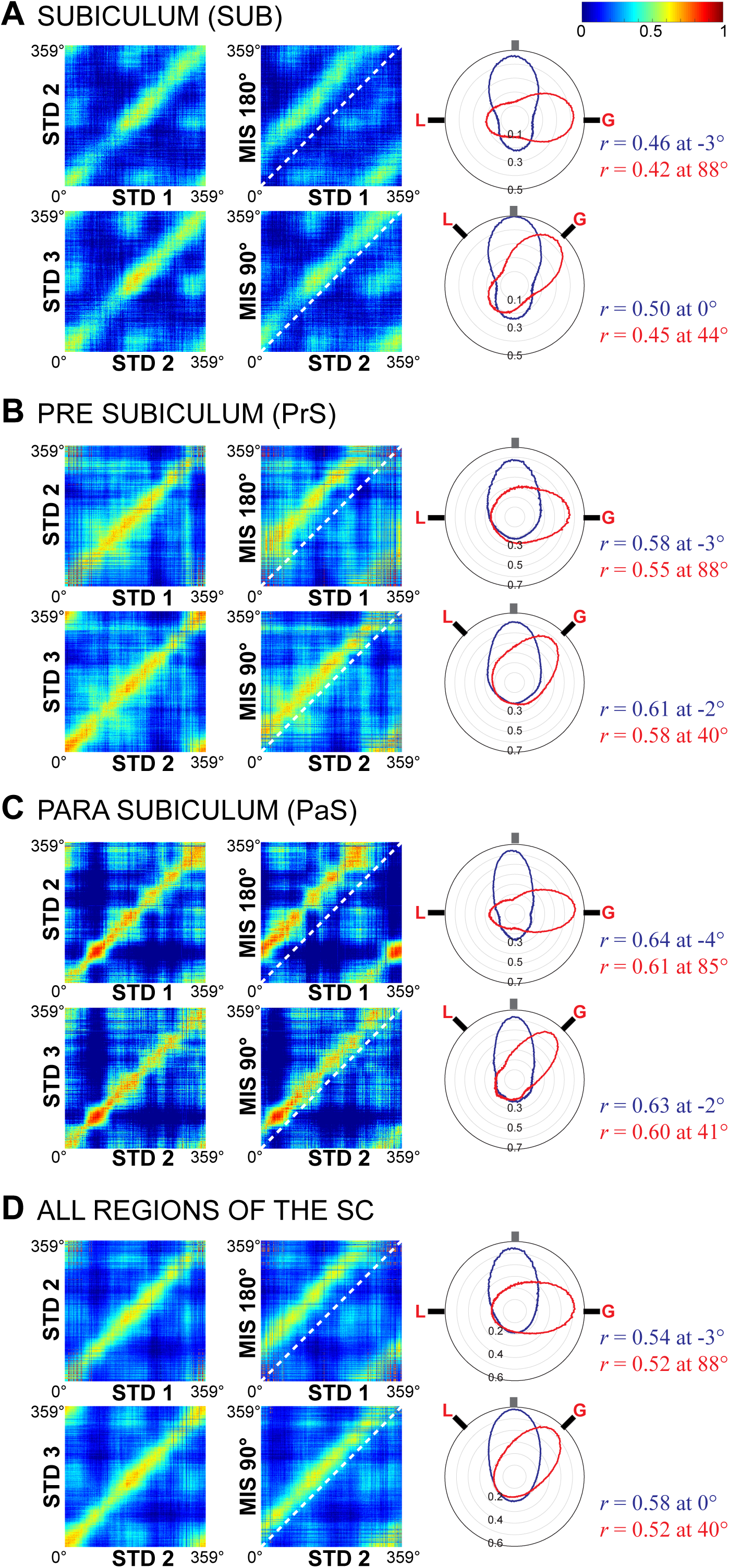
Population response in the SC network. **(A-D)** Correlation matrices from population firing rate vectors at each location on the track between STD vs. STD and STD vs. MIS sessions, created separately for the cells recorded from SUB **(A)**, PrS **(B)**, PaS **(C)** and by pooling all the cells recorded from different regions of the SC **(D).** Central diagonal is indicated by a dashed white line. The population activity between STD vs STD sessions (blue) and STD vs MIS sessions (red) are represented as polar plots (created from 2D spatial correlation matrices) and the values next to the polar plot indicates the angle at which the peak correlation (*r*) was observed. Neural representations in each of the SC regions were strongly governed by the global cues in local-global cue conflict conditions.

As we observed a highly coherent representation of the environment in the SC, much similar to an attractor-like network activity reported for the HD system, we analysed the data further to check for the possibility of coupling between the HD cells and the spatial cells in the SC by comparing the spatial offset between the co-recorded cell pairs across sessions on the circular track (*18*). Spatial cross-correlation (SXC) values were calculated for each of the co-recorded cell pairs (HD-HD, HD-Spatial) and session-wise SXC matrices were created by rank ordering the cell pairs based on their ascending peak correlation angle values in STD 1 session (Fig. 5 A and C). We observed alignment of SXC peak values along the diagonal in all session-wise SXC matrices (MIS 180°, STD 2, MIS 90° and STD 3) as the spatial offset between co-recorded cell pairs was maintained in all the sessions, indicating a unified response by all the cell pairs across sessions. As expected, the mean direction of SXCs of HD-HD cell pairs (representing the spatial offset between co-recorded HD cells within a session) across sessions (STD vs. STD, STD vs. MIS) were significantly correlated (Fig. 5 B) indicating attractor-like activity in the HD cells of the SC. Interestingly, the mean direction of SXCs of even the HD-Spatial cell pairs were also found to be significantly correlated (Fig. 5 D), suggesting a strong coupling between the HD cells and the spatial cells in the SC. The same was observed even when all the HD-Spatial cell pairs from respective STD sessions were grouped together and compared with the cue mismatch sessions (Fig. 5 E-F), indicating attractor dynamics in the SC network.

**Figure 5.**
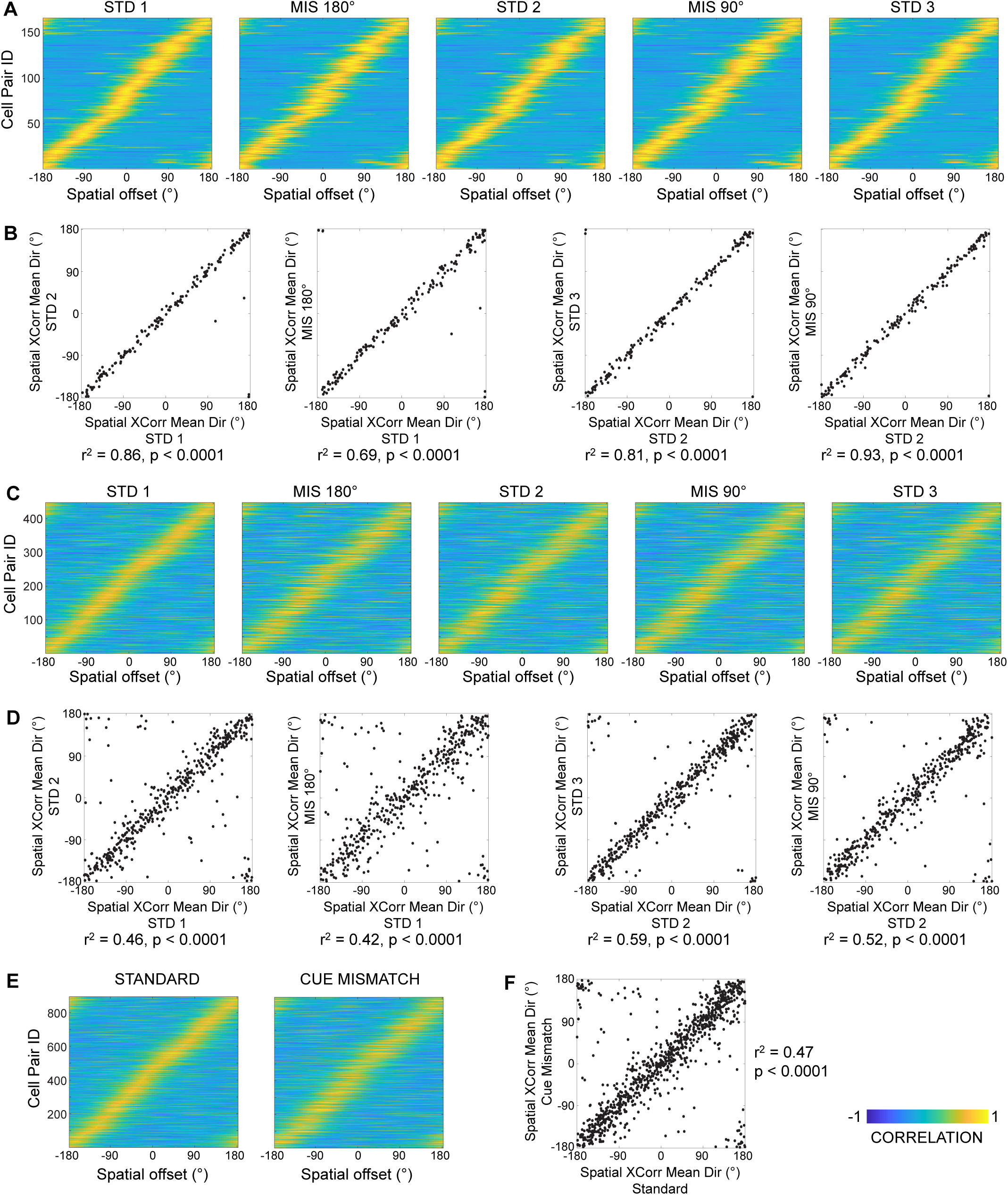
Coupling between HD cells and Spatial cells in the SC. **(A-D)** SXC matrices of co-recorded HD-HD cell pairs (167 cell pairs) **(A)** and HD-Spatial cell pairs (447 cell pairs) **(C)** across three STD and two MIS sessions, sorted based on their ascending peak correlation angle values in STD 1 session. Scatter plots showing correlation between the mean direction of SXCs of HD-HD cell pairs **(B)** and HD-Spatial cell pairs **(D)** in STD vs. STD and STD vs. MIS sessions. The *r*^*2*^ values and the significance level (*p*) are mentioned below the scatter plots for each of the comparisons. **(E-F)** Combined SXC matrices of co-recorded HD-Spatial cell pairs (894 cell pairs) and the correlation between the mean direction of SXCs of cell pairs in STD vs. Cue Mismatch sessions.

## Discussion

As per the cognitive map theory of O’Keefe and Nadel, the hippocampus is proposed to form the locus of cognitive map of the environment (*19*). The cognitive map by itself is a very complex process derived from integration of the inputs from different neural networks, involving sensory inputs, heading direction, path integration and behavioural contingencies of the animal (*20*). Various studies have reported that the spatial representation in the hippocampal circuitry is heterogeneous and performs pattern completion or pattern separation functions, because of which it can distinguish environments and create distinct cognitive maps for each of the environments based on the behavioural context and episodic events (*13-15, 17, 21, 22*). The SC receives dense projections from the hippocampal CA1 region (*23*) and SUB has been suggested to act as a major output structure of the hippocampal circuit because of its projections to various brain regions (*24*). We observed highly coherent representation of the cue-conflict environment in the SC as compared to the hippocampal representations reported under the same experimental conditions (*13-15*). Our findings clearly rule out the possibility of simple transfer of processed information from the hippocampus to other brain regions via the SC. The SC also receives topographically organized projections from the EC (*24*) and a dissociation in the representation has been reported for the EC, where the global reference frame controls the representation in the medial EC (MEC) and the local reference frame controls the representation in the lateral EC (LEC) (*16*). It is unlikely that the SC may be reflecting only the MEC representations as the coherency of representations in the SC appears much stronger than the global or local cue controlled representations in the MEC or LEC respectively. Moreover, our observations of robust coupling between HD cells and Spatial cells of the SC shows compelling similarity with the attractor-like representation in the ADN HD network (*10, 15, 18*).

It has been suggested that the ADN-HD system may drive the neural activity in dorsal part of the PrS, the postsubiculum (PoS) (*25*) and shows dependence on the PoS for landmark cue control (*26*) which is critical for path integration and navigation in familiar environments. The PoS has strong reciprocal connections with the ADN (*27, 28*) and lesioning of the PoS leads to loss of visual landmark cue control over ADN HD cells (*26*), suggesting an integration of the external visual information on to the incoming HD signal at the PoS region. Moreover, the PoS HD cells have also been shown to carry spatial information (*29*), and developmental studies in rats have reported generation of an adult like HD signal in PoS and MEC neurons much prior to spatial information processing activity (*30, 31*). Interestingly, the coherency of representation in our data was not limited only to the PoS HD cells, but was observed in all sub-regions of the SC and in different cell types of the SC. The intrinsic connectivity amongst SUB, PrS and PaS along with reciprocal connections with the HD system (*1*) may endow the SC to act as an attractor network, resulting in strong coupling between the HD and spatial cells, and a highly coherent representation of the environment.

Even though different sub-regions of the SC network may perform distinct functions, such as the role of SUB in memory retrieval (*32*), PrS for landmark orientation (*26*) and PaS in representing online spatial information (*33*), the intrinsic anatomical connections amongst these regions may enable SC to develop a generic framework by encoding the prominent cues of an environment and provide a consistent spatio-directional code for orientation to other interdependent neural circuits to form a specific spatial representation of the environment. In other words, the neural computations in the SC may lead to formation of a ‘reference map’ of an environment, in line with the notion of a ‘universal map’ in the SUB proposed by Sharp (*34, 35*). This ‘reference map’ may be thought of as a template that must constantly update itself from animal’s heading direction to provide correct orientation and the selection of frame of reference for orientation may be governed by the cues that are perceived to be stable through rapid re-weighting based on the cue reliability (*36*). Any failure in the generation or updating of which may lead to cognitively dysfunctional spatial representation. For example, lesioning of the SUB leading to deficits in heading and landmark bearing (*37*) or impaired pattern separation process in the hippocampus (*38*), lesioning of the PoS disrupting stability of the hippocampal place cells in familiar environment (*39*), and combined lesioning of the PoS and the PaS affecting the spatial tuning of place cells (*40*) leading to impaired spatial navigation, spatial memory and contextual conditioning (*41-43*).

We conclude that our data showing characteristic features of the attractor dynamics, a highly coherent spatial representation along with robust coupling between the HD cells and Spatial cells in the SC, is the first empirical evidence to demonstrate the unique role of the SC network in providing a common spatial platform in the form of a ‘reference map’ of the environment, which might then be relied upon by the EC and the hippocampus to form a cognitive map of the environment, along with further processing in terms of behavioral contingencies, context and episodic events.

## Materials and Methods

### Subjects and surgical procedures

Long-Evans rats (n = 5, male, 5-6 months old) were housed individually on reversed light-dark (12:12 h) cycle and the experiments were carried out during the dark phase of the cycle. All the procedures (animal care, surgical procedures and euthanasia) were performed in accordance with NIH guidelines and were approved by the Institutional Animal Ethics Committee (IAEC) of National Brain Research Centre at Manesar, Haryana, constituted by the Committee for the Purpose of Control and Supervision of Experiments on Animals (CPCSEA), Government of India. The surgical procedures were performed under aseptic conditions. Initially, the rats were anaesthetised using ketamine (60 mg/kg b.w.) and xylazine (8mg/kg b.w.) and then shifted to isoflurane gaseous anaesthesia for the entire duration of the surgery. A custom-built recording device (Microdrive) containing 20 independently movable tetrodes (18 tetrodes and 2 references, contained inside a single bundle or dual bundle) was surgically implanted over the right hemisphere in a way to simultaneously access different regions of the subicular complex (SC) at 5.5 - 8.5 mm posterior to bregma and 2.5 – 4.5 mm lateral to midline. Post-surgical care was provided upon surgical implantation of the microdrive, and the rats were allowed to recover over a period of 7 days. Following post-surgical recovery, the tetrodes were lowered into the brain targeting different regions of the SC and the rats were trained to run clockwise on a circular track. During training and subsequent experimental recordings, the rats were maintained at 85% of their free feeding weights.

### Electrophysiology and recording

Tetrodes were made from 17 μm platinum-iridium wire (California Fine Wire, USA) and the tips of individual wires were electroplated with platinum black solution (Neuralynx Inc., USA) to 100-150 kΩ with 0.2 μA current. Electrophysiological recordings were carried out using 96-channel data acquisition system (Digital Lynx 10S, Neuralynx Inc., USA) by amplifying the signals through headstage preamplifier (Neuralynx Inc., USA). The units were recorded against a reference electrode present in a cell-free zone in the brain by filtering the signal between 600 Hz and 6 kHz. Spike waveforms above the threshold were sampled for 1 ms at 32 kHz. Local field potentials were recorded against a ground screw anchored to the skull above frontal cortex, filtered between 0.1 Hz and 1 kHz, and continuously sampled at 4 kHz. The position and the head direction of the animal was tracked with the red and green LEDs attached to the headstages, captured through ceiling mounted color CCD camera (CV-S3200, JAI Inc, San Jose, USA) at 25 Hz.

### Behavior and recording procedures

After 7 days post-surgical recovery, the tetrodes were slowly advanced targeting different regions of the SC over a period of 10-15 days, by keeping the rat on a pedestal next to the recording system. During this period, the rats were trained in the adjacent behavioral room to run clockwise seeking chocolate sprinkles placed at random locations on a centrally placed textured circular track (elevated 90 cm from floor level) for 30 min/day for 8-10 days. The circular track (56 cm inner diameter; 76 cm outer diameter) with four types of visually distinct textured surfaces, each covering a quarter on the track, served as local cues. Six salient cues (four visually distinct cues of different pattern and shape made of cardboard hung on curtain, two cardboard boxes of different size placed on floor at the periphery of curtain) at the surrounding black curtain reaching from ceiling to the floor, served as global cues. A ceiling mounted circular LED light at the centre of the room provided required illumination and a centrally placed white noise generator on the floor masked any external sounds during training and subsequent experimental sessions. Once tetrodes reached the region of interest and the animal was trained, the experimental recording sessions were carried out as described below in 5 rats, following the experimental protocol designed by Prof. James J. Knierim (*13*).

### Local–Global cue conflict experiment

Each day of recording consisted of five sessions, in which three standard sessions (STD) were interleaved with two mismatch sessions (MIS). In STD sessions, the configuration between the local cues and the global cues was maintained exactly same as that during the training sessions. Local – Global cue conflict was created in MIS sessions by rotating all the global cues clockwise (CW) and the textured circular track counter clockwise (CCW) by either 90° or 45° each to create a mismatch of 180° (MIS-180°) or 90° (MIS-90°) between the cues.

### Experimental procedure

The rat was first placed inside a covered box for 30 seconds, and carried by the experimenter on a brief walk in the computer room and around the track inside the behavioral room to disrupt rat’s sense of direction between the external environment and the behavioral room. A pedestal was placed in the centre of the textured track, the rat was transferred onto the pedestal, recording tethers with headstages was connected to the electrode interface board of the microdrive, the rat was released onto the track at random starting point and the pedestal was removed. Upon completion of 20 laps, the pedestal was returned to the centre of the track, the rat was placed on the pedestal to disconnect the tethers, transferred to the box and taken on a brief walk. This procedure was repeated for each of the experimental sessions. On completion of all the experimental sessions, the track was wiped clean with 70% ethanol to clear off any traces that could act as potential cues for next day of recording. In order to classify different types of the SC cells being recorded, each day after the last session on the track, a circular platform (76 cm diameter) was placed on top of the textured track and the neural activity was recorded for 10 min duration while the rat foraged in the arena.

## Data analysis

The analysis of data was performed with custom-written software, MATLAB (R2013a) and circular statistics (Oriana, Kovach Computing Services, UK), as described below.

### Isolation of single-units

Isolation of single-units was performed manually with custom-written spike-sorting software Winclust, developed by Prof. James Knierim, Johns Hopkins University, Baltimore, USA. Cells were isolated based on the peak amplitude and energy of the waveforms recorded on four wires of the tetrode. Based on their isolation quality (distance from the background and separation from other clusters), the units were rated on a scale ranging from 1 to 5 (1-very good; 2-good; 3-fair; 4-marginal; 5-poor) and the units rated ‘fair’ and above were used for further analysis.

### Cell type identification

Different types of cells have been reported in the SC. The cells recorded were classified as Head direction cell, Border cell, Grid cell or Place cell based on their Rayleigh’s mean vector length or Border score or Grid score values that were above the threshold values obtained through data shuffling. These scores were calculated from the recordings on a circular platform at the end of track session.

To calculate border score and grid score, spatial firing rate maps were created for all the cells recorded by binning the recording area into 64×48 bins (bin size approx. 2×2 cm). The number of spikes in each bin was divided by the amount of time spent in that bin to obtain a bin-wise firing rate matrix of individual cells, which was then smoothed using a Gaussian kernel of 5×5 bins with a sigma of 2. To calculate Rayleigh’s mean vector length, head directional firing rate tuning curves were created for all the cells recorded by dividing the number of spikes fired when rat’s head was facing a particular direction (bin size 5°) by the total amount of time spent facing that particular direction.

#### Border score

The MATLAB codes for calculating the border score was obtained through personal communications from Prof. Edward Moser’s laboratory, and the border score was computed based on the formula as described in Solstad et. al., 2008 (*44*);

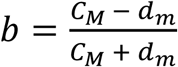

where, *b* is the border score computed for each cell, *C*_*M*_ is the maximum coverage of any single field over any of the boundary of the environment, *d*_*m*_ is the mean firing distance computed by averaging the distance from any of the fields to the nearest boundary over all pixels in the map, weighted by the firing rate. A cell was classified as border cell if its border score was more than the 99^th^ percentile of border score values (threshold value) obtained through data shuffling as described in Langston et. al., 2010 (*31*) (see below for the data shuffling procedure).

#### Grid score

Grid score was calculated by using the MATLAB codes available online (https://github.com/derdikman/Ismakov-et-al.-Matlab-code) (*45*). 2D autocorrelograms were generated from spatial firing rate maps of each cell by calculating Pearson’s product moment correlation between the original rate map of a cell with its shifted version having spatial lags of τ_x_ and τ_y_:

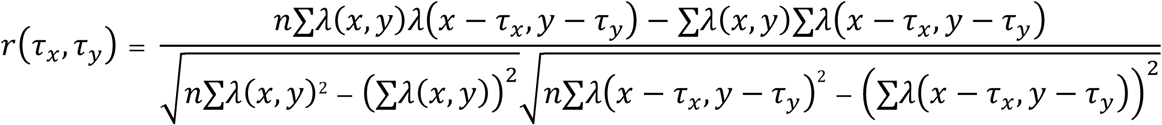

where *λ* (*x, y*) denotes the average rates of a cell at location (*x,y*), the autocorrelation between these fields with spatial lags of *τ* _*x*_ and *τ* _*y*_ was estimated. The summation is over all *n* pixels in *λ* (*x, y*) for which rate was estimated for both *λ* (*x, y*) and *λ* (*x* −*τ* _*x*_, *y* −*τ* _*y*_) (*46*).

Gridness of a cell was defined as the periodicity of the peaks in the spatial 2D-autocorrelogram. The autocorrelogram was correlated with its rotated version (excluding the central peak) for every angle centered around its central peak and Pearson’s correlation value for 30°, 60°, 90°, 120° and 150° rotation was obtained. Grid cells have a repeated pattern of peaks to give high correlation values at 60° and 120° and lower correlation values at 30°, 90° and 150°. Gridness score is the difference between the minimum correlation value at 60° or 120° and maximum correlation value at 30° or 90° or 150° (*47*). A cell was defined as a grid cell if its grid score was more than the 99^th^ percentile of grid score value obtained by data shuffling (*31*) (see below for the data shuffling procedure).

#### Rayleigh’s mean vector length

Rayleigh’s mean vector length is the measure of directional modulation of firing rate of a cell across all head directions. The mean vector length for a cell with higher firing rates along narrow range of head directions is higher as compared to a cell with firing rates spread along all head directions. The Rayleigh’s vector was calculated in MATLAB by following circular statistical analysis (*48*), where the mean firing rate in every angular bin_i_ was considered as a point on a unit circle with X_j_ and Y_j_ coordinates, using the formula;

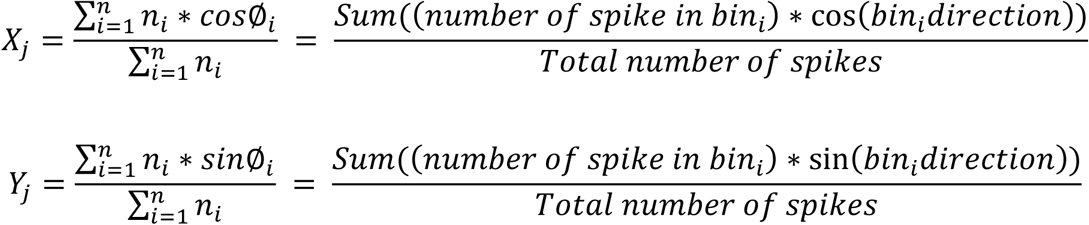

where, values in each angular bin_i_ constitutes one vector pointing to a particular direction. The sum of all the individual vectors divided by the total number of vector N giving the mean resultant vector 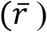, whose absolute value is the Rayleigh vector (R):

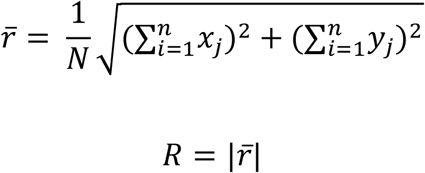

A cell was classified as a Head direction cell if its Rayleigh’s vector score value was more than the 99^th^ percentile of Rayleigh’s vector score values obtained through data shuffling (see below for data shuffling procedure), provided its border/grid score value was less than those corresponding threshold values.

#### Data Shuffling

Data shuffling was performed to rule out the possibility of spuriously classifying the cells by chance to different types. Shuffling was performed separately for each cell where a cell’s entire spike sequence recorded on the platform session was time shifted by a random interval between 20^th^ second and 20 seconds before termination of the session. Spikes exceeding the total time of the session were wrapped around to be assigned to the beginning of session to generate new spike time sequence for the cell. This procedure was repeated 100 times for each cell, and the Border score, Grid score and Rayleigh’s mean vector length were calculated on these shuffled spike time series. 99^th^ percentile value of shuffled distribution of each score was taken as the threshold value for that cell type (*31*).

#### Spatial information score

The spatial information score was calculated for all those remaining cells that were not categorised as either a border cell, grid cell or head direction cell. The spatial information value indicates the amount of information about the rat’s position, conveyed by the firing of a single spike from a cell (*49*), and was calculated as;

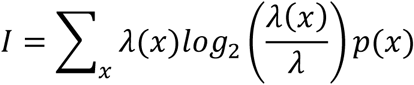

where x is spatial bin, λ(x) is the firing rate of the cell at location, λ is the mean firing rate and p(x) is the probability of occupancy at bin x. A cell was classified as a Place cell if its spatial information score was found to be significant (*p* < 0.05) in any of the track sessions recorded in a day.

### Rotational correlation

In order to assess the change in preferred firing direction/location of the HD cells and Spatial cells (place, border and grid cells), we performed the rotational correlation analysis by following the methods described in Neunuebel et. al., 2013 (*16*). To begin with, two-dimensional circular track data was linearized using 5° binning to obtain one dimensional firing rate array (of 72 bins) for each cell. Directional tuning curve was calculated for head-direction cells by dividing total number of spikes fired when animal’s head faced a particular direction by the total amount of time spent facing that particular direction on the circular track. In case of Spatial cells, one dimensional firing rate arrays were created by dividing the number of spikes fired when the animal was in a particular spatial bin by the total amount of time spent in that particular bin on the circular track. A speed filter of 1cm/sec was used for all Spatial cells to avoid accumulation of spikes fired by a cell when the animal was too slow or stationary on the circular track. To quantify the amount of rotation of cells between STD sessions or between STD and MIS sessions, we have measured the Pearson’s product-moment correlation of linearized firing rate arrays of a cell between those two sessions. Then, the firing rate bins of the session being correlated was shifted by one bin (5° shift) to obtain shifted correlations, and this procedure was repeated for 71 times. The shift angle at which maximum correlation value was obtained, was considered as the amount of rotation of a cell between the two sessions being compared. We performed circular statistical tests (*50*) to calculate the angle of the mean vector and the length of the mean vector, using circular statistics software (Oriana, Kovach Computing Services, UK). The mean angle of rotation of the cells was represented by the angle of the vector, while the length of the vector was inversely proportional to the variance of the distribution around that mean. The angle of the mean vector for all CW rotations were between 0° to 180°, while the same for CCW rotations were between 0° to −180°.

### Population coherence

To understand the SC network activity, the population responses were measured by creating 2D spatial correlation matrices from population firing rate vectors at each location on the track. Population correlation matrices for STD vs STD sessions and STD vs MIS session were created for all the cells recorded. Firing rate vectors for each session were created by pooling firing rate arrays (bin size 1°) of all the cells from a particular session yielding a N x 360 matrix where N is the number of cells. The standard session firing rate vector at each bin was correlated (using Pearson’s product-moment correlation) with subsequent standard or mismatch session bin to produce a 360 x 360 correlation coefficient matrix. A band of high correlation in the population correlation matrix signified coherent activity where, band along the diagonal signified no rotation. Bands above or below the diagonal represents CW or CCW rotations of cell population, as illustrated in figure below. In order to quantify the population response, the 2D spatial correlation matrices were reduced to 1D polar plots by calculating the mean correlation of pixels in each of the 360 diagonals in the population correlation matrices. The angle at which maximum mean correlations were obtained was considered as the amount of rotation of the cell population between the two sessions (*14, 16, 51*).

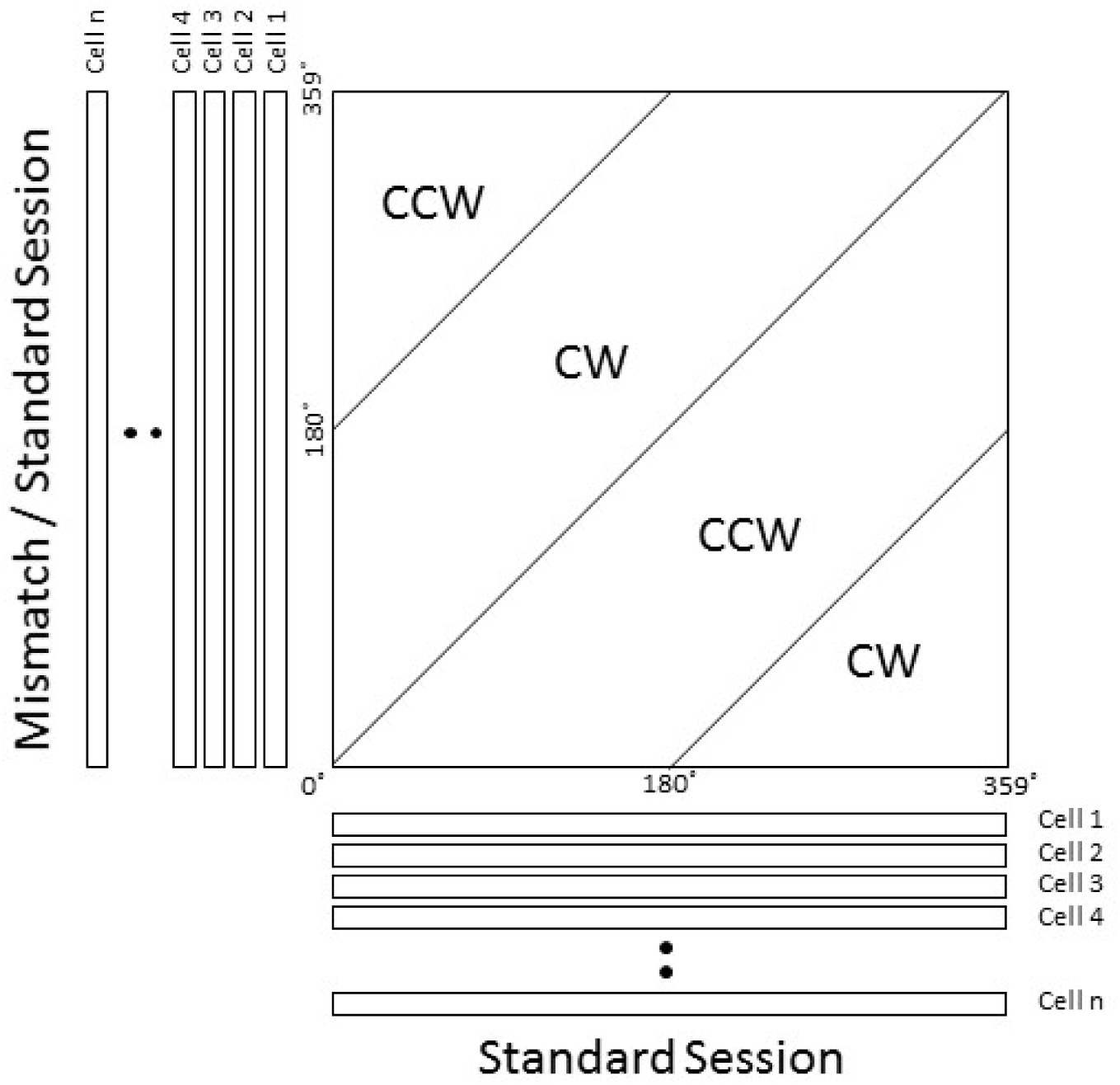

### Spatial cross-correlation (SXC)

To assess the possibility of coupling between HD cells and Spatial cells in the SC, the spatial offset between the co-recorded cell pairs across sessions on the circular track was compared by adapting the procedure described in Bassett et al 2018 (*18*), with an exception that the spatial cross correlations were performed on the spike data for the whole session, as illustrated in the schematic below;

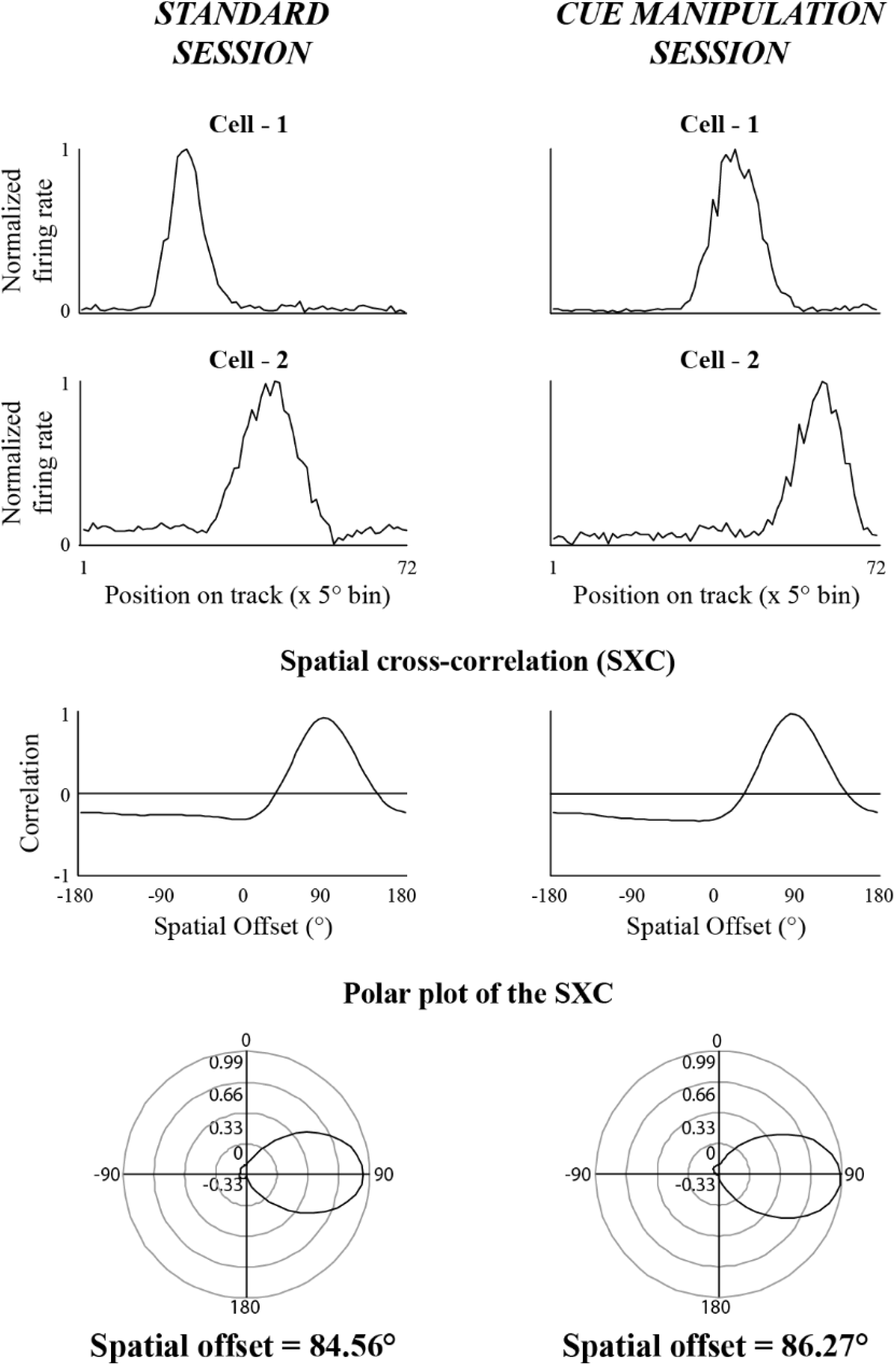

One dimensional firing rate array (of 72 bins) for each cell was created as described above under rotational correlation analysis, and were normalized to its peak firing rate. The SXC values were calculated through Pearson’s product-moment correlation of normalized firing rate arrays of a cell pair in a session, and the normalized firing rate array of the cell being compared was shifted by one bin (5° shift) to obtain shifted correlation value, and the procedure was repeated for 71 times. The angle at which maximum SXC was observed for a cell pair was defined as the peak correlation angle value. SXC matrices were created by pooling the cell pairs from all days of recordings, session-wise, and rank ordered based on its ascending peak correlation angle value in STD 1 session for visualizing the alignment of SXC peak values across various sessions. For quantitative analysis, polar plots of SXCs were created for each of the cell pairs and mean direction of the polar plot was calculated using circular statistics toolbox (*52*) in MATLAB. The mean direction value represented the spatial offset between co-recorded cells within a session on the circular track. Through correlation analysis, the coupling between the cells were quantified by comparing the spatial offset values of cell pairs across sessions, and the r-squared values (indicating goodness of fit of the linear equation) and its significance were calculated. As the HD system is proposed to exhibit attractor dynamics (*53-55*), we first assessed the coupling between co-recorded HD-HD cell pairs across sessions in datasets having a minimum of two HD cells. Upon confirmation of an attractor-like coupling amongst the HD cells in the SC, we quantified the coupling between co-recorded HD-Spatial cell pairs across sessions.

#### Histological procedures and identification of recording sites

Upon completion of the electrophysiological experiments, marker lesions were performed on few selected tetrodes by passing current (10 μA for 10 seconds). On the following day, rats were perfused transcardially with 4% formalin solution, the brain was extracted, and stored in 30% sucrose-formalin until it sank in the solution. Brains were sectioned in the coronal plane (40 μm thick), mounted, and processed for Nissil’s staining using 0.1% Cresyl violet. Images of serial sections were captured on Leica DFC265 digital camera attached to Leica M165-C stereo microscope and saved as TIFF files. The distance from midline to the tetrode track markings were measured from these serial sections, plotted in excel spreadsheet to visualize the configuration of tetrode tracks. The tetrodes were identified by comparing this configuration with the arrangement of tetrodes in the microdrive, and cross verifying with the marker lesions. We performed depth reconstruction of the tetrode track to identify the brain region at which the cells were recorded on each day, based on the distance from the bottom tip of the tetrode by taking into account a 15% shrinkage of tissue due to histological processing.

## References

1. S. L. Ding, Comparative anatomy of the prosubiculum, subiculum, presubiculum, postsubiculum, and parasubiculum in human, monkey, and rodent. J. Comp. Neurol. 521, 4145–4162 (2013).

2. C. A. Barnes, B. L. McNaughton, S. J. Mizumori, B. W. Leonard, L. H. Lin, Comparison of spatial and temporal characteristics of neuronal activity in sequential stages of hippocampal processing. Prog. Brain Res. 83, 287–300 (1990).

3. P. E. Sharp, C. Green, Spatial correlates of firing patterns of single cells in the subiculum of the freely moving rat. J. Neurosci. 14, 2339–2356 (1994).

4. P. E. Sharp, Multiple spatial/behavioral correlates for cells in the rat postsubiculum: multiple regression analysis and comparison to other hippocampal areas. Cereb. Cortex 6(2), 238–259 (1996).

5. J. S. Taube, Place cells recorded in the parasubiculum of freely moving rats. Hippocampus 5, 569–583 (1995).

6. J. S. Taube, R. U. Muller, J. B. Jr. Ranck, Head-direction cells recorded from the postsubiculum in freely moving rats. I. Description and quantitative analysis. J. Neurosci. 10, 420–435 (1990).

7. C. N. Boccara, F. Sargolini, V. H. Thoresen, T. Solstad, M. P. Witter, E. I. Moser, M. B. Moser, Grid cells in pre- and parasubiculum. Nat. Neurosci. 13, 987–994 (2010).

8. C. Lever, S. Burton, A. Jeewajee, J. O’Keefe, N. Burgess, Boundary Vector Cells in the Subiculum of the Hippocampal Formation. J. Neurosci. 29, 9771–9777 (2009).

9. J. S. Taube, R. U. Muller, J. B. Jr. Ranck, Head-direction cells recorded from the postsubiculum in freely moving rats. II. Effects of environmental manipulations. J. Neurosci. 10(2), 436–447 (1990).

10. J. S. Taube, H. L. Burton, Head direction cell activity monitored in a novel environment and during a cue conflict situation. J. Neurophysiol. 74, 1953–1971 (1995).

11. P. E. Sharp, Subicular cells generate similar spatial firing patterns in two geometrically and visually distinctive environments: comparison with hippocampal place cells. Behav. Brain Res. 85, 71–92 (1997).

12. J. R. Brotons-Mas, N. Montejo, S. M. O’Mara, M. V. Sanchez-Vives, Stability of subicular place fields across multiple light and dark transitions. Eur. J. Neurosci. 32(4), 648–658 (2010).

13. J. J. Knierim, Dynamic interactions between local surface cues, distal landmarks, and intrinsic circuitry in hippocampal place cells. J. Neurosci. 22, 6254–6264 (2002).

14. I. Lee, D. Yoganarasimha, G. Rao, J. J. Knierim, Comparison of population coherence of place cells in hippocampal subfields CA1 and CA3. Nature 430, 456–459 (2004).

15. D. Yoganarasimha, X. Yu, J. J. Knierim, Head direction cell representations maintain internal coherence during conflicting proximal and distal cue rotations: comparison with hippocampal place cells. J. Neurosci. 26, 622–631 (2006).

16. J. P. Neunuebel, D. Yoganarasimha, G. Rao, J. J. Knierim, Conflicts between Local and Global Spatial Frameworks Dissociate Neural Representations of the Lateral and Medial Entorhinal Cortex. J. Neurosci. 33, 9246–9258 (2013).

17. J. P. Neunuebel, J. J. Knierim, CA3 Retrieves Coherent Representations from Degraded Input: Direct Evidence for CA3 Pattern Completion and Dentate Gyrus Pattern Separation. Neuron 81, 416–427 (2014).

18. J. P. Bassett, T. J. Wills, F. Cacucci, Self-Organized Attractor Dynamics in the Developing Head Direction Circuit. Current Biology 28, 609–615 (2018).

19. J. O’Keefe, L. Nadel, The hippocampus as a cognitive map. Clarendon, Oxford, (1978).

20. L. F. Jacobs, F. Schenk, Unpacking the cognitive map: The parallel map theory of hippocampal function. Psychological Review, 110(2), 285–315 (2003).

21. T. J. Wills, Attractor Dynamics in the Hippocampal Representation of the Local Environment. Science, 308, 873–876 (2005).

22. J. K. Leutgeb, S. Leutgeb, M. B. Moser, E. I. Moser, Pattern Separation in the Dentate Gyrus and CA3 of the Hippocampus. Science, 315, 961–966 (2007).

23. D. G. Amaral, C. Dolorfo, P. Alvarez-Royo, Organization of CA1 projections to the subiculum: a PHA-L analysis in the rat. Hippocampus 1(4), 415–435 (1991).

24. M. P. Witter, Connections of the subiculum of the rat: topography in relation to columnar and laminar organization. Behav. Brain Res. 174, 251–264 (2006).

25. A. Peyrache, M. M. Lacroix, P. C. Petersen, G. Buzsáki, Internally organized mechanisms of the head direction sense. Nature Neuroscience 18, 569–575 (2015).

26. J. P. Goodridge, J. S. Taube, Interaction between the postsubiculum and anterior thalamus in the generation of head direction cell activity. J. Neurosci. 17, 9315–9330 (1997).

27. T. van Groen, J. M. Wyss, The postsubicular cortex in the rat: characterization of the fourth region of the subicular cortex and its connections. Brain Res. 529, 165–177 (1990).

28. T. van Groen, J. M. Wyss, Projections from the anterodorsal and anteroventral nucleus of the thalamus to the limbic cortex in the rat. J. Comp. Neurol. 358, 584–604 (1995).

29. A. Peyrache, N. Schieferstein, G. Buzsaki, Transformation of the head-direction signal into a spatial code. Nat. Commun. 8, 1752 (2017).

30. T. J. Wills, F. Cacucci, N. Burgess, J. O’Keefe, Development of the hippocampal cognitive map in preweanling rats. Science 328, 1573–1576 (2010).

31. R. F. Langston, J. A. Ainge, J. J. Couey, C. B. Canto, T. L. Bjerknes, M. P. Witter, E. I. Moser, M. B. Moser, Development of the spatial representation system in the rat. Science 328, 1576–1580 (2010).

32. D. S. Roy, T. Kitamura, T. Okuyama, S. K. Ogawa, C. Sun, Y. Obata, A. Yoshiki, S. Tonegawa, Distinct Neural Circuits for the Formation and Retrieval of Episodic Memories. Cell 170(5), 1000–1012 (2017).

33. Q. Tang, A. Burgalossi, C. L. Ebbesen, J. I. Sanguinetti-Scheck, H. Schmidt, J. J. Tukker, R. Naumann, S. Ray, P. Preston-Ferrer, D. Schmitz, M. Brecht, Functional Architecture of the Rat Parasubiculum. J. Neurosci. 36(7), 2289–2301 (2016).

34. P. E. Sharp, Complimentary Roles for Hippocampal Versus Subicular/Entorhinal Place Cells in Coding Place, Context, and Events. Hippocampus 9, 432–443 (1999).

35. P. E. Sharp, Subicular place cells generate the same “map” for different environments: comparison with hippocampal cells. Behav Brain Res. 174(2), 206-214 (2006).

36. K. J. Jeffery, H. J. I. Page, S. M. Stringer, Optimal cue combination and landmark-stability learning in the head direction system. J. Physiol. 594(22), 6527–6534 (2016).

37. R. G. M. Morris, F. Schenk, F. Tweedie, L. E. Jarrard. (1990). Ibotenate Lesions of Hippocampus and/or Subiculum: Dissociating Components of Allocentric Spatial Learning. Eur. J. Neurosci. 2(12), 1016–1028 (1990).

38. O. Potvin, F. Y. Doré, S. Goulet, Lesions of the dorsal subiculum and the dorsal hippocampus impaired pattern separation in a task using distinct and overlapping visual stimuli. Neurobiol Learn Mem. 91(3), 287–297 (2009).

39. J. L. Calton, R. W. Stackman, J. P. Goodridge, W. B. Archey, P. A. Dudchenko, J. S. Taube, Hippocampal place cell instability after lesions of the head direction cell network. J Neurosci. 23, 9719–9731 (2003).

40. P. Liu, L. E. Jarrard, D. K. Bilkey, Excitotoxic lesions of the pre- and parasubiculum disrupt the place fields of hippocampal pyramidal cells. Hippocampus 14, 107–116 (2004).

41. J. S. Taube, J. P. Kesslak, C. W. Cotman, Lesions of the rat postsubiculum impair performance on spatial tasks. Behav. Neural Biol. 57, 131–143 (1992).

42. R. P. Kesner, R. Giles, Neural circuit analysis of spatial working memory: role of pre- and parasubiculum, medial and lateral entorhinal cortex. Hippocampus 8, 416 – 423 (1998).

43. P. Liu, L. E. Jarrard, D. K. Bilkey, Excitotoxic lesions of the pre- and parasubiculum disrupt object recognition and spatial memory processes. Behav. Neurosci. 115, 112–124 (2001).

44. T. Solstad, C. Boccara, E. Kropff, M. B. Moser, E. I. Moser, Representation of geometric borders in the entorhinal cortex. Science 322, 1865–1868 (2008).

45. R. Ismakov, O. Barak, K. Jeffery, D. Derdikman, Grid Cells Encode Local Positional Information. Current Biology 27(15), 2337–2343 (2017).

46. T. Hafting, M. Fyhn, S. Molden, M. B. Moser, E. I. Moser, Microstructure of a spatial map in the entorhinal cortex. Nature 436, 801–806 (2005).

47. F. Sargolini, M. Fyhn, T. Hafting, B. L. McNaughton, M. P. Witter, M. B. Moser, E. I. Moser, Conjunctive representation of position, direction and velocity in entorhinal cortex. Science 312, 758–762 (2006).

48. N. I. Fisher, Statistical Analysis of Circular Data. Cambridge University Press (1995).

49. W. E. Skaggs, B. L. McNaughton, K. M. Gothard, E. J. Markus, An information-theoretic approach to deciphering the hippocampal code. Adv. Neural Inf. Process Syst. 5, 1030–1037 (1993).

50. J. H. Zar, Biostatistical Analysis, 4th edition (Prentice Hall, Upper Saddle River, NJ, 1999).

51. K. M. Gothard, W. E. Skaggs, B. L. McNaughton, Dynamics of mismatch correction in the hippocampal ensemble code for space: interaction between path integration and environmental cues. J. Neurosci. 16, 8027–8040 (1996).

52. P. Berens, CircStat: A MATLAB Toolbox for Circular Statistics. J. Statistical Software 31, 1–21 (2009).

53. W. E. Skaggs, J. J. Knierim, H. Kudrimoti, B. L. McNaughton, A model of the neural basis of the rat’s sense of direction. Neural Information Processing Systems 7, 173–180 (1995).

54. K. Zhang, Representation of spatial orientation by the intrinsic dynamics of the head-direction cell ensemble: a theory. J. Neurosci. 16, 2112–2126 (1996).

55. A. D. Redish, A. N. Elga, D. S. Touretzky, A coupled attractor model of the rodent head direction system. Network: Computation in Neural Systems 7, 671–685 (1996).

